# Individual variation in parental care drives divergence of sex roles

**DOI:** 10.1101/2020.10.18.344218

**Authors:** Xiaoyan Long, Franz J. Weissing

## Abstract

In many animal species, parents provide care for their offspring, but the parental roles of the two sexes differ considerably between and within species. Here, we use an individual-based simulation approach to investigate the evolutionary emergence and stability of parental roles. Our conclusions are in striking contrast to the results of analytical models. In the absence of initial differences between the sexes, our simulations do not predict the evolution of egalitarian care, but either female-biased or male-biased care. When the sexes differ in their pre-mating investment, the sex with the highest investment tends to evolve a higher level of parental care; this outcome does not depend on non-random mating or uncertainty of paternity. If parental investment evolves jointly with sexual selection strategies, evolution results in either the combination of female-biased care and female choosiness or in male-biased care and the absence of female preferences. The simulations suggest that the parental care pattern drives sexual selection, and not *vice versa*. Finally, our model reveals that a population can rapidly switch from one type of equilibrium to another one, suggesting that parental sex roles are evolutionarily labile. By combining simulation results with fitness calculations, we argue that all these results are caused by the emergence of individual variation in parental care strategies, a factor that was hitherto largely neglected in sex-role evolution theory.

## Introduction

In the animal kingdom, species differ remarkably in the way and degree female and male parents are involved in parental care^1,2^. In virtually all mammals, most of the care is provided by females^1,3^, while in birds biparental care (with a certain bias towards females) is the most prevalent pattern^1,4^. Teleost fishes exhibit a broad variety of care patterns, with male-biased care being the rule rather than the exception^1,5^. Even within species, parental care patterns can be highly diverse^6^. For example, in Eurasian penduline tits (*Remiz pendulinus*) female-only care and male-only care co-occur in the same population^7^, while in Chinese penduline tits (*Remiz consobrinus*) female-only care, male-only care, and biparental care all coexist^8^. Moreover, phylogenetic studies suggest that parental care patterns are highly dynamic in that transitions between patterns occur frequently^9,10,11^.

The explanations that have been proposed for sex differences in parental roles often initiated heated debates in the literature. One debate centres around the role of anisogamy (the difference in gamete size between males and females). Robert Trivers^12^ argued that anisogamy explains the fact that in many taxa females tend to invest more in post-zygotic parental care than males. According to Trivers, the female parent has a strategic disadvantage with respect to the male parent: because the mother has made a large initial investment in the ovum, she has more to lose when deserting the clutch than the father. Some authors pointed out a flaw in Trivers’ argument: optimal decision-making should not be based on past investments, but rather on future costs and benefits^13^. While agreeing with this critique, other authors pointed out that Trivers’ prediction can be revived when taking other factors into account, such as female choosiness or uncertainty of paternity^14,15^. This viewpoint is, in turn, hotly debated^16–19^. Another debate in the literature is on whether and how the relative abundance of males and females drives parental sex roles^20^. A popular theory predicts that the ‘operational sex ratio’ (the ratio of males to females among those individuals participating in mating) should play a decisive role, because the sex that is overrepresented on the mating market (and hence has fewer mating opportunities) should be predestined for taking on the parental care tasks^21^. More recently, attention has shifted to the ‘adult sex ratio’ (the ratio of males to females in the overall adult population) as a predictor of sex differences in parental sex roles^22,23^. Last, but not least, there is debate in the literature on the role of sexual selection in determining parental sex roles^12,14,24^. All these debates are intricate in themselves; moreover, they are interwoven, because initial investments, sex ratios, and sexual selection are mutually dependent^12^.

In a situation like this, where the outcome of evolution is determined by the intricate interplay of mutually dependent factors, verbal theories can easily lead astray. As a major step forward, Kokko and Jennions^25^ developed a comprehensive modelling framework, allowing to disentangle the role of the various factors involved in the evolution of parental sex roles. In a first step, male and female fitness functions are calculated, based on a scheme describing the interactions of the sexes in a population. These functions are then analysed mathematically (see Methods), allowing to predict how sex differences in life history parameters, biased sex ratios, multiple mating, and sexual selection affect the evolution of parental sex roles. However, this analytical approach has its limitations. First, the calculations are not trivial and error-prone. Indeed, Fromhage and Jennions^26^ pointed out mistakes and erroneous conclusions in the study of Kokko and Jennions^25^. Second, to keep the model analytically tractable, the factors involved have to be stripped to their bare-bone essentials. For example, the dynamic process of sexual selection is reduced to a set of fixed parameters that cannot coevolve with the parental strategies. Third, the analytical approach focuses on the evolution of population means and thereby neglects intra-population variation around the mean. In other words, populations are considered monomorphic, while it has recently become clear that in natural populations individuals differ systematically in all kinds of behavioural tendencies^27,28,29^, including parental behaviour^30,31,32^. Various studies have shown that such variation is often shaped by diversifying selection^33,34^, and that it can have important evolutionary implications^35,36^.

For these reasons, we here consider an extended version of the modelling framework of Kokko and Jennions^25^, and we study the evolution of parental roles by means of individual-based simulations^37^.

This approach has the advantage that more natural assumptions can be made concerning the inclusion of sexual selection or factors such as sex differences in pre-mating investment. Moreover, individual variation emerges in a natural way, making it possible to study its evolutionary implications^37^.

In a nutshell, our model (see Methods) follows individual males and females from birth to death. After maturation, adult individuals can be in one of two states: the mating state and the caring state. Individuals seek mating opportunities in the mating state; once mated both members of the mated pair switch to the caring state. Each individual provides care for a time period corresponding to its inherited sex-specific parental care strategy and switches back to the mating state afterwards. The total amount of care provided by both parents determines the survival probability of the offspring in the clutch. The offspring inherit the care strategies from their parents (according to Mendelian inheritance and subject to rare mutations of small effect size). Parental care strategies have to strike a balance between caring as efficiently as possible and mating as often as possible. Both caring and mating are costly, since individuals can die in any state, with a mortality rate that depends on their state and sex. Strategies that perform well are transmitted to a large number of offspring, thereby increasing in relative frequency in the population. Over the generations, an evolutionary equilibrium emerges during the simulation; fitness calculations are not required for this. As explained below, the model can easily be extended to include sexual selection and sex-differences in pre-mating investment.

Although the model is very similar in spirit to the analytical models mentioned above, we show that the evolutionary outcome is remarkably different from that reported in the earlier studies of parental sex-role evolution.

## Results

### Sex-biased care evolves in the absence of sex differences

First, we consider the baseline scenario where mating is at random and the sexes do not differ in their mortality rates or other life-history parameters (Fig. 1). Based on their analytical model, Kokko and Jennions^25^ predicted the evolution of egalitarian biparental care for this scenario. Correcting a mistake in the fitness calculations, Fromhage and Jennions^26^ showed that instead the analytical model predicts convergence to a line of equilibria. If we apply the selection gradient method of refs 25 and 26 (see Supplementary Fig. 5 - 7) to our slightly modified model, we arrive at the same conclusion (Fig. 1a): the care effort of females and males converges to an equilibrium; there is a continuum of equilibria, which are located on a curve that includes a broad spectrum of parental care patterns. In other words, depending on the initial conditions all types of care strategy, from female-only care via egalitarian biparental care to male-only care, can evolve.

**Fig. 1.**
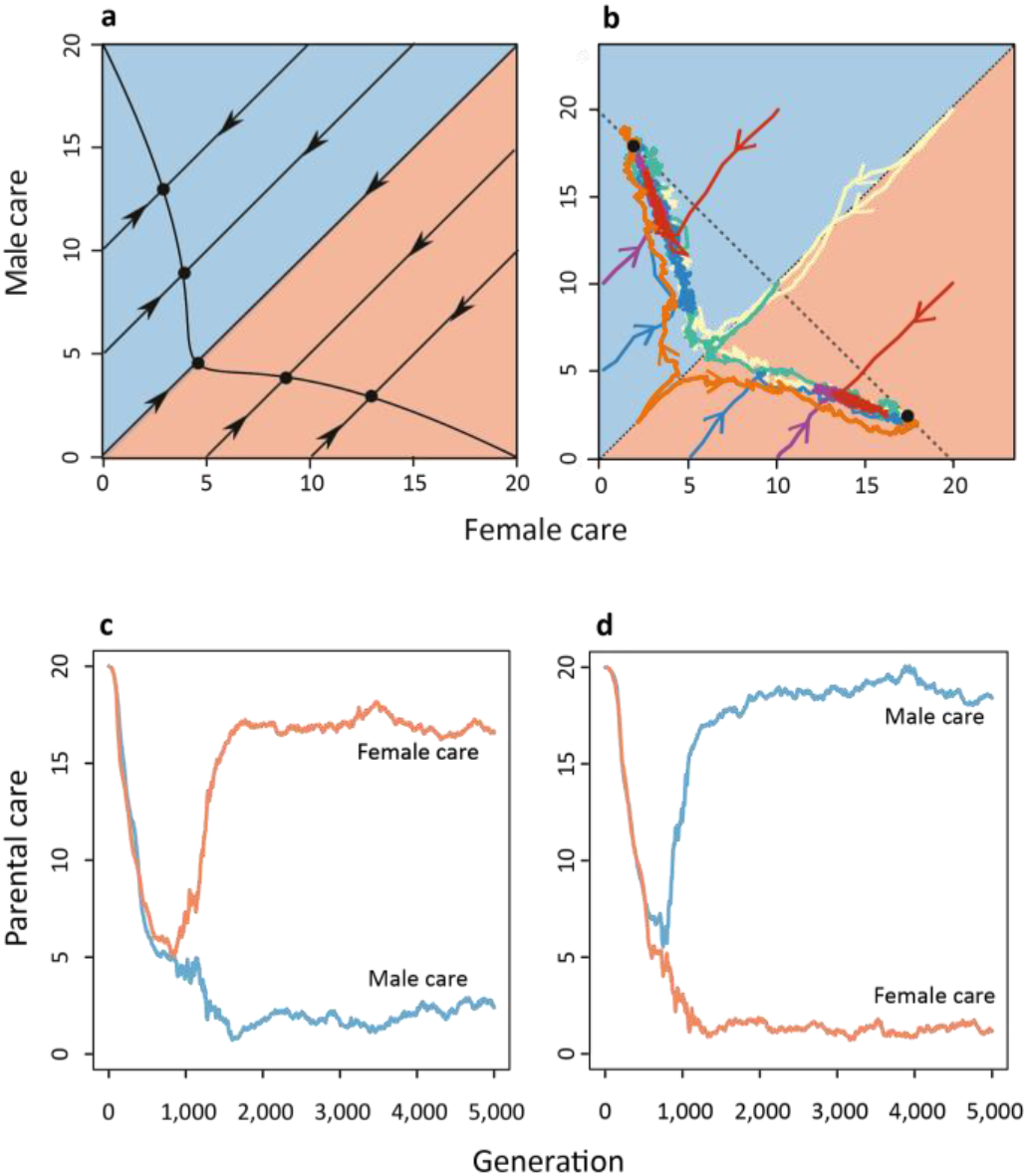
Evolution of sex-biased parental roles in the absence of sexual selection. The graphs depict evolutionary trajectories when mating is at random and males and females do not differ in mortality rates or other life-history parameters. **(a)** For this scenario, the selection gradient method predicts convergence to a curve of equilibria (solid black line). **(b)** In contrast, individual-based simulations converge in a characteristic manner to one of two equilibria (black dots) corresponding to either strongly female-biased care or strongly male-biased care. Replicate simulations starting with egalitarian care levels will converge, with equal probability, to **(c)** the female-care equilibrium or **(d)** the male-care equilibrium. Differently coloured lines in (b) indicate different initial conditions. The red and blue lines in (c) and (d) depict the average levels of female care and male care in the evolving population. The dotted line in (b) corresponds to those care levels where the sum of female and male care is equal to *D* = 20, the value of total care maximizing the marginal benefits of care in our model (see Methods). Population sizes fluctuated around 2,000 females and 2,000 males.

In contrast to these analytical predictions, our individual-based simulations never resulted in egalitarian care or a line (or curve) of equilibria. Instead, all our simulations (>5,000, for different parameter values and different initial conditions) converged to one of two stable equilibria corresponding to either strongly female-biased care or strongly male-biased care. Initial conditions with sex-biased care tended to converge to the corresponding sex-biased equilibrium, while initial conditions without sex-bias converged to each of the two equilibria with equal probability (Fig. 1b). Fig. 1c and 1d show the time trajectories of two replicate simulations starting at a high level of egalitarian care. In a first phase, both populations follow the analytical prediction and converge to a low level of egalitarian care. Then strongly sex-biased care evolves, along the curve of equilibria of the analytical model. Both stable equilibria have the property that the total care provided by the two parents equals *D* = 20, the value maximizing the marginal benefit of care in our model (see Methods).

### The evolution of sex-biased parental roles is driven by individual variation

Fig. 2 shows in more detail how sex-biased care evolves from egalitarian care. In the simulation shown, the population was initialized at the same care level (20) for males and females. Hence, initially the sum of the parental care levels exceeds the value *D* = 20 that, for the parameters chosen, maximizes the marginal benefits of care. Accordingly, there is strong selection in both sexes to reduce the level of care. In the first 800 generations, the care level in males and females rapidly declines until a value of 5 is reached in both sexes (Fig. 2a,b), in line with the predictions of the selection gradient approach (see Fig. 1a). At this care level, the mortality of offspring is very high and additional care would provide a considerable benefit. Yet, the parents are caught in a cooperation dilemma: both are interested in the survival of their offspring, but each parent is better off if most of the care is provided by the other parent^12,38,39^

**Fig. 2.**
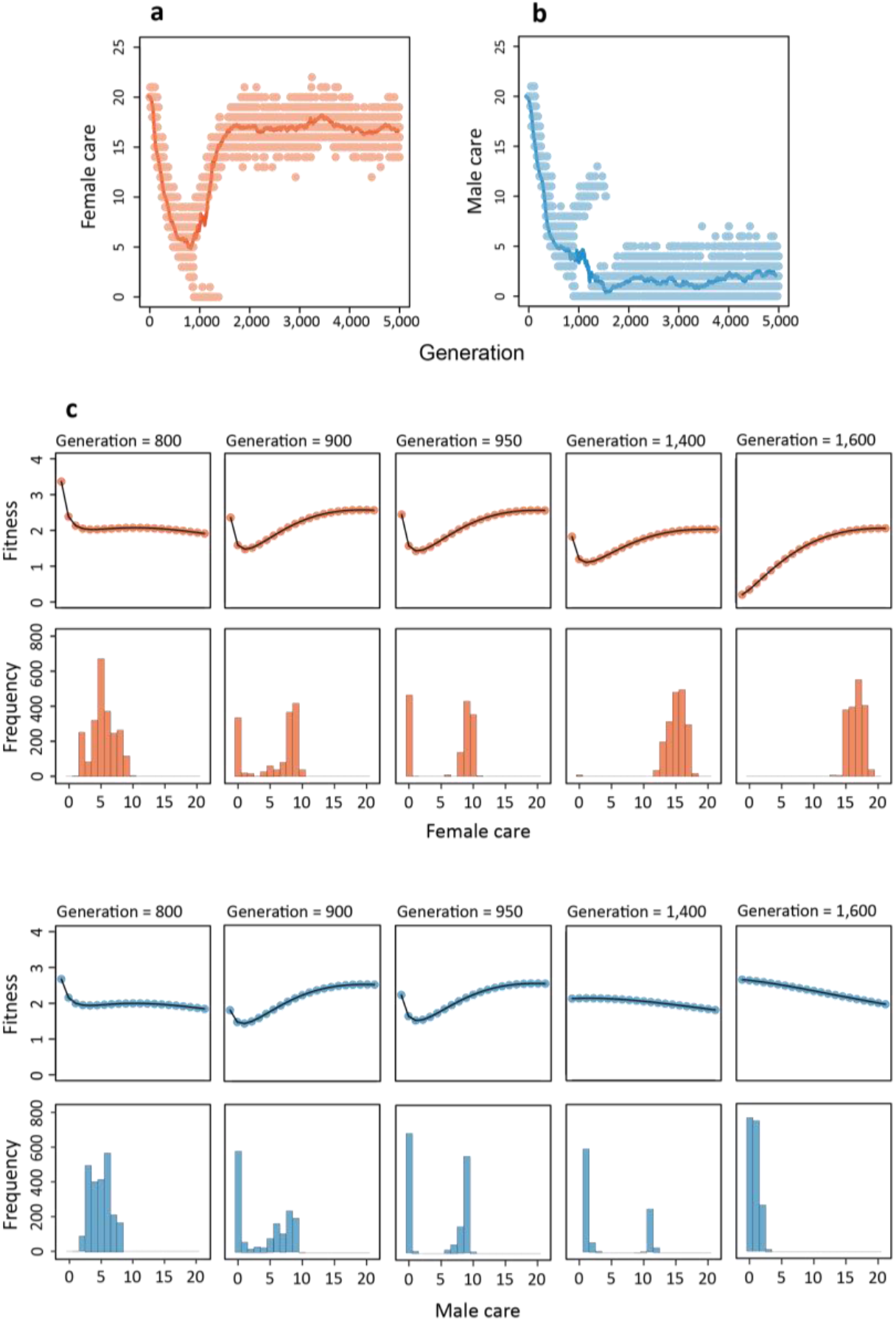
Sex role divergence driven by individual variation in parental roles. Evolution of **(a)** female and **(b)** male care for the simulation in Fig. 1c. Lines show the average care level of females (red) and males (blue) in the population, while dots represent individual care levels. **(c)** For five different generations, the histograms show the distribution of care levels in females (red) and males (blue). The fitness profiles above the histograms indicate in each case the expected lifetime reproductive success of females and males with care strategies ranging from 0 to 20 in the corresponding population.

To understand the further course of evolution, we first considered the simplified version of the model where parental care is constrained to be egalitarian (i.e., individuals cannot determine their care duration dependent on their sex). In this egalitarian model, a care level of 5 for both parents corresponds to an ‘evolutionary branching point’^40^ (see Supplementary Fig. 2): at such a point, directional selection changes into disruptive selection, where extreme strategies have the highest fitness. This is confirmed by the U-shaped fitness profile and the emerging bimodal distribution of care levels in both sexes in generation 900 (see Fig. 2c). The process continues, and in generation 950 there are two types of females and two types of males: one type not caring at all and the other type caring at a level around 10. In the egalitarian version of the model, the process would continue until part of the population would not care at all while the other part would care at level *D* = 20. Such a population is not very efficient, because many matings would result in either no care at all or a very high care level of 40. When individuals can make their care strategy dependent on their sex (or any other phenotypic marker), there is an escape route^41^: one of the two ‘branches’ becomes associated with the female sex, while the other becomes associated with the male sex. In the simulation in Fig. 2, the high-care strategy becomes associated with the female sex and the no-care strategy becomes associated with the male (the opposite happened in 50% of the simulations): in generation 1400, the no-care strategy has almost disappeared in females and selection is directional in males (in favour of the no-care strategy). In the end (generation 1600), directional selection keeps the care level low in males, while stabilizing selection keeps the care level just below 20 in females. Without exception, the same sequence of events (with similar timing) was observed in hundreds of simulations starting with similar care levels in the two sexes.

### Anisogamy affects the evolution of parental sex roles even in the absence of sexual selection

In most taxa females tend to invest more in post-zygotic parental care than males^1–4^. Since females are, by definition, the sex producing larger gametes, it is plausible to assume that anisogamy plays an important role in the evolution of parental sex roles^18,26^. Trivers’ argument that the sex with the highest pre-mating investment is predestined to invest more in post-zygotic care because it has ‘more to lose’ is generally considered to be flawed^13^, but various authors pointed out other causal links from anisogamy to female-biased care, via secondary effects of anisogamy, such as higher competition among males or a lower certainty of parentage in males^14,15^. To investigate the role of pre-mating investment, we extended our model by introducing a pre-mating period for one of the sexes. After any parental care period, an individual of that sex has to spend a fixed number of days with other activities (like growing a new clutch of eggs in females or building a new nest in males) before entering the mating phase again. Mating is still assumed to be at random, and there are no other differences between the sexes.

Fig. 3 shows, for two mortality levels in the pre-mating period, that the sex with the higher pre-mating investment tends to evolve a higher degree of post-zygotic parental care. This trend is very pronounced (white curve) if the mortality in the pre-mating period is five times as high as in the mating period, but it is also noticeable when the pre-mating period does not involve direct fitness costs, because the mortality level is zero (black curve). Hence, we clearly observe a ‘Trivers effect’ in the absence of sexual selection and multiple matings. We think that this outcome results from the interplay of two factors. First, a longer pre-mating period leads to a shorter life expectancy, which shifts the balance between current and future reproduction toward a higher investment in the current clutch^42,43^. Second, the sex with the shorter pre-mating period has a higher variance in mating success, which selects for higher mating effort and reduced parental care^44^. The first factor does not play a role when there is no mortality in the pre-mating period (because in that case life expectancy is not affected). Fig. 3 (black dots and line) demonstrates that even in that case the second factor, which was first predicted by Sutherland^44^, has a noticeable effect on the evolutionary outcome. In other words, Trivers was right, but for different reasons than he envisaged. Additional implications of anisogamy, such as paternity uncertainty or intrinsically more intense competition among males are not required but will most probably enhance the Trivers effect.

**Fig. 3.**
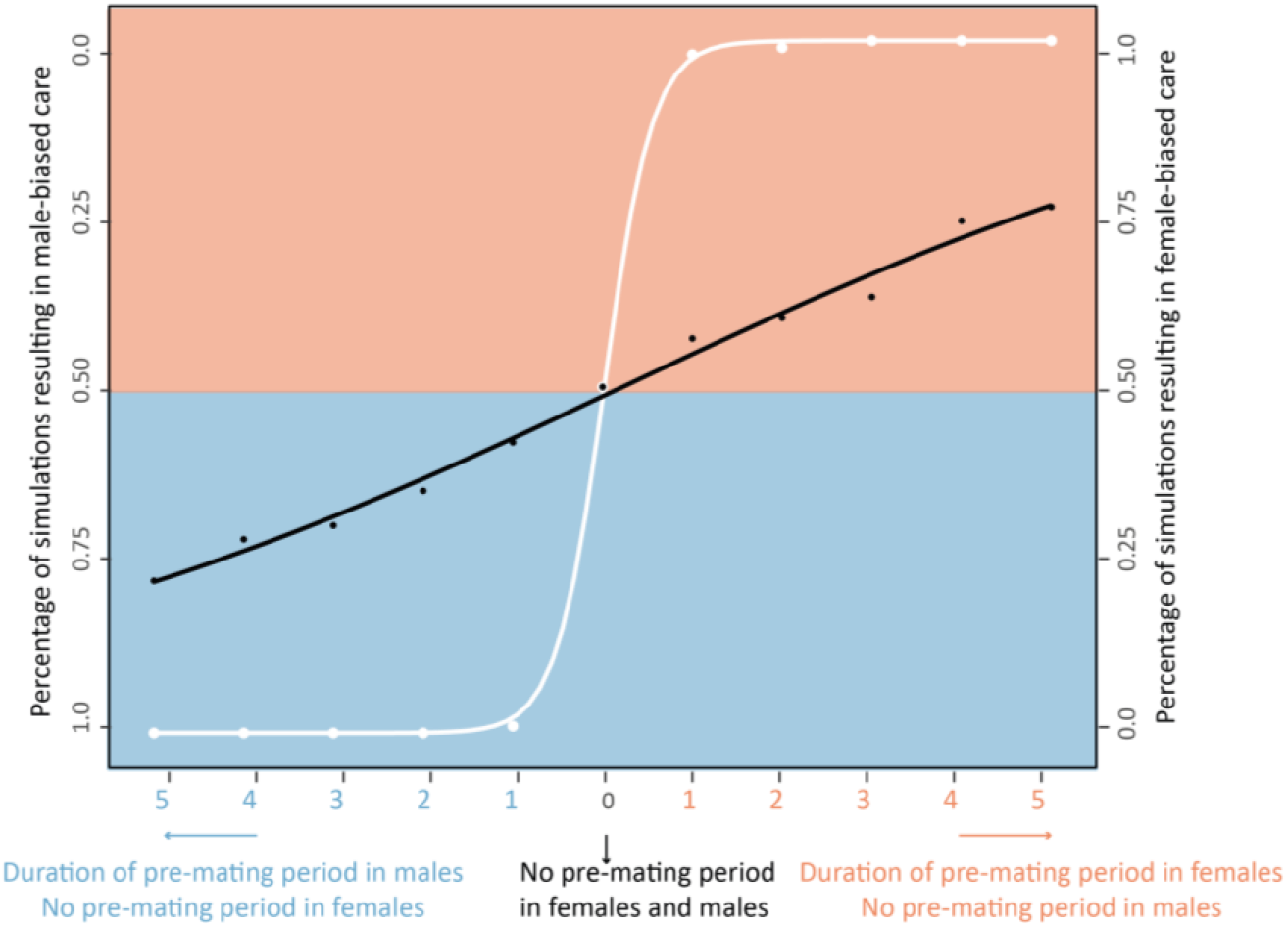
A pre-mating investment bias selects for parental sex roles. Percentage of simulations resulting in male-biased care (left axis) or female-based care (right axis) depending on the duration of the pre-mating period in either males (blue) or females (red). Mortality in the pre-mating period was either zero (black dots and black line fitted by logistic model) or five times as high as in the mating phase (white dots and white line fitted by logistic model). 100 replicate simulations were run per setting, all starting from egalitarian care. All of these 2,200 simulations resulted either in female-biased care or male-biased care. In case of a female pre-mating period, female-biased care was the more likely outcome, while male-biased care evolved more often when males had a pre-mating period.

### Parental sex roles can be evolutionarily labile

Up to now, all simulations converged to one of two alternative equilibria that correspond to either male-biased or female-biased care. As shown in Fig. 4, rapid switches from one equilibrium to the other were regularly observed on a long-term perspective. In fact, such switches *always* occurred in situations with alternative stable equilibria, provided that the simulations were run for a sufficiently long time period. Accordingly, our simulations suggest that parental roles can be evolutionarily labile. This is in line with phylogenetic studies, which also conclude that parental care patterns are highly dynamic and that, on a long-term perspective, transitions between different care patterns have occurred frequently in many animal taxa^9,10,11^.

**Figure 4.**
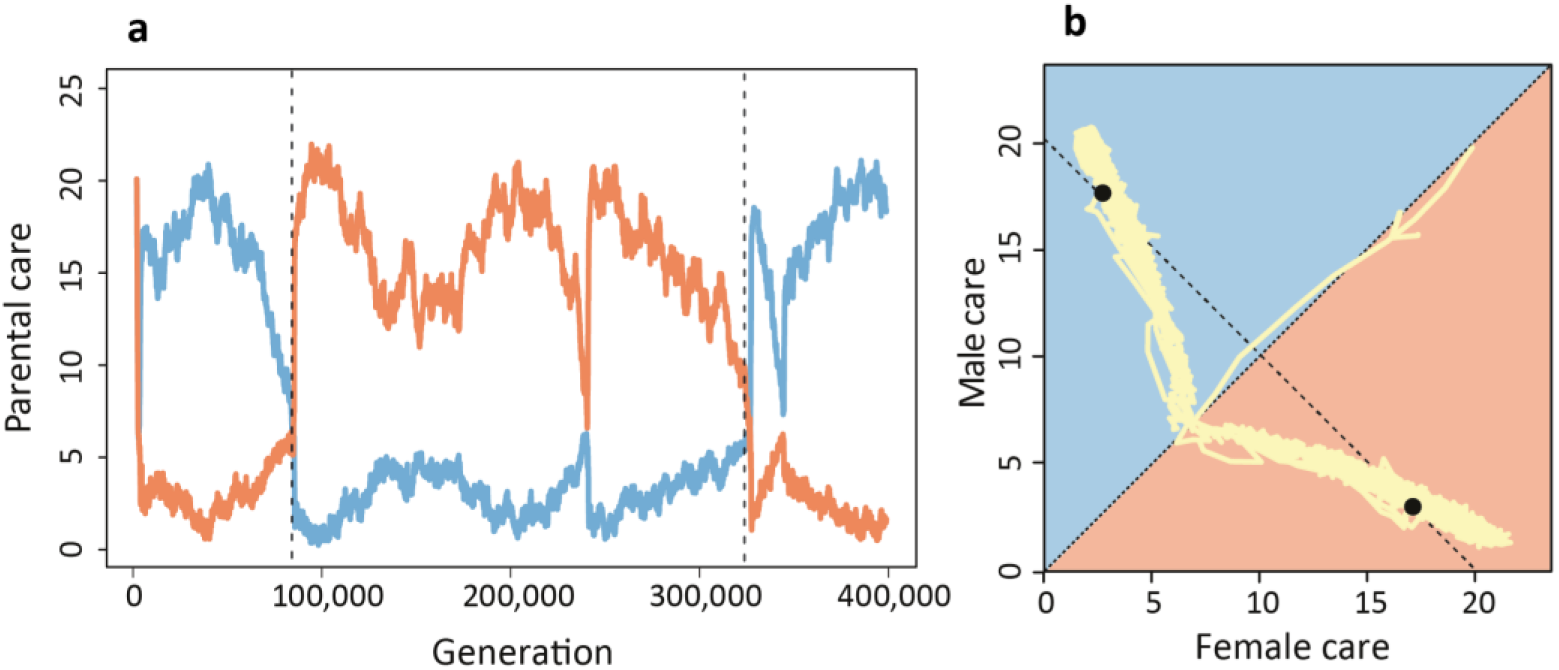
Evolutionary lability of parental sex roles. Whenever simulations were run for extended periods of time, transitions occurred between the two stable equilibria. In other words, long periods of male- or female-biased care were followed by rapid switches to a situation where most of the care was provided by the other sex. Here, this is shown for a long-term simulation of the random-mating scenario in Fig. 1, but with a one-day pre-mating period in both sexes.

In a stochastic dynamical system with alternative stable states, spontaneous transitions from one state to the other are not really surprising^45^. They occur, for example, in ecological systems^46,47^, in the climate system^48^, and in physical systems^49^ (think of the spontaneous reversal of polarity in magnets^50^). The average time between switches depends on the degree of stochasticity and the strength of attraction, which in our case corresponds to population size and the steepness of the selection gradients. Decreasing the population size by relaxing density dependence did indeed lead to much faster transitions between states (see Supplementary Fig. 3). The same happened when we weakened selection by prolonging the pre-mating period in one or both sexes (as in Fig. 4).

### Biparental synergy can lead to fluctuating polymorphism or inefficient biparental care

In contrast to the simulations reported above, egalitarian biparental care occurs in many bird and fish species, and in other animal taxa^1–4^. A potential reason is that in natural populations the parents complement each other, thereby providing more benefits to their offspring than the sum of their individual contributions^51^. Division of labour or other sources of synergy among the parents could reduce sexual conflict about who should do the caring and strongly select for biparental care^52,53^. Here we introduce parental synergy in our model in line with earlier modelling studies^26,52^: we assume that the care levels *T*_*f*_ and *T*_*m*_ of the two parents provide a benefit *T*_*f*_ + *T*_*m*_ + *σT*_*f*_*T*_*m*_ to their offspring, where the degree of synergy *σ* is a positive parameter (In the additive model considered until now, *σ* = 0). In the analytical model of Fromhage and Jennions^26^, the introduction of a small degree of synergy transformed their curve of equilibria (Supplementary Fig. 5) into a single stable equilibrium corresponding to egalitarian biparental care.

Fig. 5 shows that this prediction is only partly confirmed by individual-based simulations. When synergy is weak (*σ* = 0.05, Fig. 5a), the population does not converge to an equilibrium. Instead, the average care level in both sexes (top panel of Fig. 5a) exhibits large fluctuations, corresponding to rapid transitions between female-biased and male-biased care. Moreover, both sexes are polymorphic most of the time: a considerable fraction of individuals does not care at all, while others provide a high level of care. In case of an intermediate degree of synergy (*σ* = 0.20, Fig. 5b), the population converges to egalitarian care, although both the male and the female population remain highly polymorphic. Notice that the average care level (top panel of Fig. 5b) in both sexes is about *T*_*f*_ = *T*_*m*_ = 5 and, hence, very low. Taking synergy into account, this investment results in a total care level of about 5 + 5 + 0.2 + 25 = 15. This is considerably less than in the additive model without synergy (Fig. 1b), where in both non-egalitarian equilibria the total care level is equal to *D* = 20, the value maximizing the marginal benefits of parental care. Apparently, the introduction of synergism does not allow the parents to escape from the cooperation dilemma by the evolution of either male-biased or female-biased care. Instead, the conflict between the sexes continues, resulting in a broad spectrum of care strategies and an outcome that is, regarding offspring survival, quite inefficient. This conclusion only changes for a high degree of synergy (*σ* = 2.0, Fig. 5c): now the population converges to an egalitarian care level satisfying *T*_*f*_ + *T*_*m*_ + *σT*_*f*_*T*_*m*_ = *D*.

**Figure 5.**
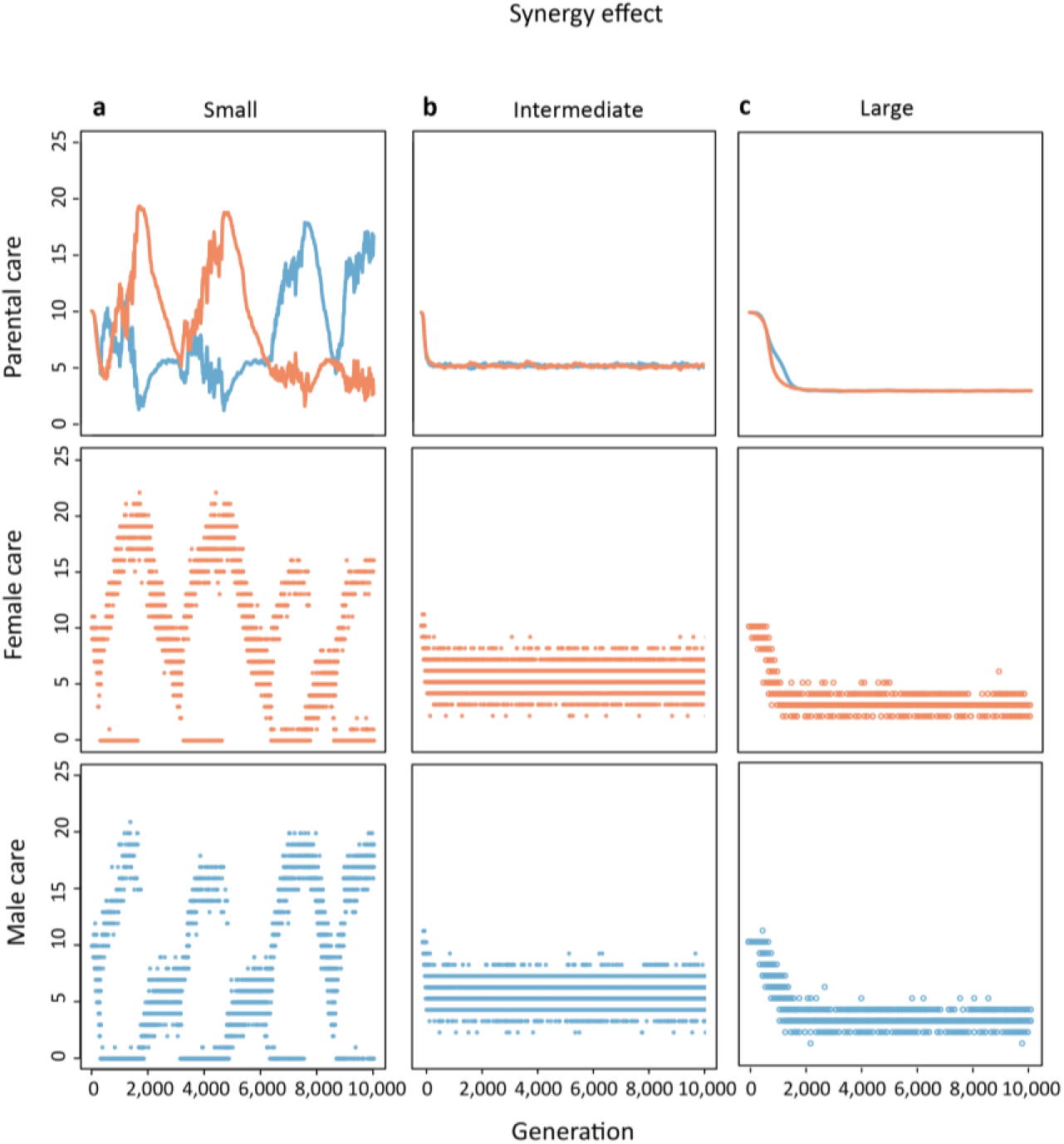
Evolution of parental roles when biparental care has a synergistic effect. Representative simulations for the case that the effects of the parents on offspring survival are not additive but synergistic. **(a)** In case of weak synergy (*σ* = 0.05), evolution leads to a rapid succession of male-and female biased care. For long periods of time, one or both sexes are highly polymorphic, with a no-care strategy coexisting with a high-care strategy. **(b)** In case of intermediate synergy (*σ* = 0.20), evolution leads to egalitarian care equilibrium. However, diverse care strategies coexist in both sexes. Total care *T*_*f*_ + *T*_*m*_ + *σT*_*f*_*T*_*m*_ is considerably smaller than *D* = 20, the value maximizing the marginal benefit of care in our model. **(c)** In case of strong synergy (*σ* = 2.0), the evolving egalitarian-care equilibrium exhibits relatively little variation and total care now matches *D* = 20.

### Joint evolution of mating and parental strategies

Mating and parental care strategies are closely interrelated, but the causal relationships between the two types of strategy are difficult to disentangle. Mathematical models incorporating both factors tend to be analytically intractable and can only be solved by iteration methods^52^. Many models on the evolution of parental roles therefore represent mating patterns by a parameter that cannot change in time^25,26^. It is a clear advantage of individual-based simulation models that various scenarios for the joint evolution of mating and parental care strategies can be implemented in a natural way. To demonstrate this, we extended the baseline version of the model by allowing female preferences and male ornaments to evolve alongside with the parental strategies. We restrict ourselves to a simple model of sexual selection, leaving the analysis of more complicated scenarios (e.g., mutual mate choice, differences in parental ability, condition-dependent mating and parental strategies) to a future attempt. In our Fisherian model^54^, female preferences and male ornaments are characterized by heritable parameters *p* and *s*, respectively. When female preferences are zero, all males have the same probability of being chosen and mating occurs at random. When female preferences are above zero, males with large ornaments are preferred. Male ornamentation is costly in that it negatively affects male survival. Female choosiness is costly, because choosy females may take a longer time before they find a mate. Fig. 6 shows some representative simulations, all starting with random mating (*p* = *s* = 0) but with different initial levels of parental care. All simulations converge to one of two equilibria (with equal probability) that are characterized by either male-biased care or female-biased care. Whenever male-biased care evolved (Fig. 6b), female preferences stayed at a very low level, corresponding to random mating. Whenever female-biased care evolved (Fig. 6c), female preferences for male ornaments evolved as well, together with elaborate male ornamentation. In all simulations leading to female-biased care, female choosiness only got off the ground after female care levels had reached relatively high levels.

**Fig. 6.**
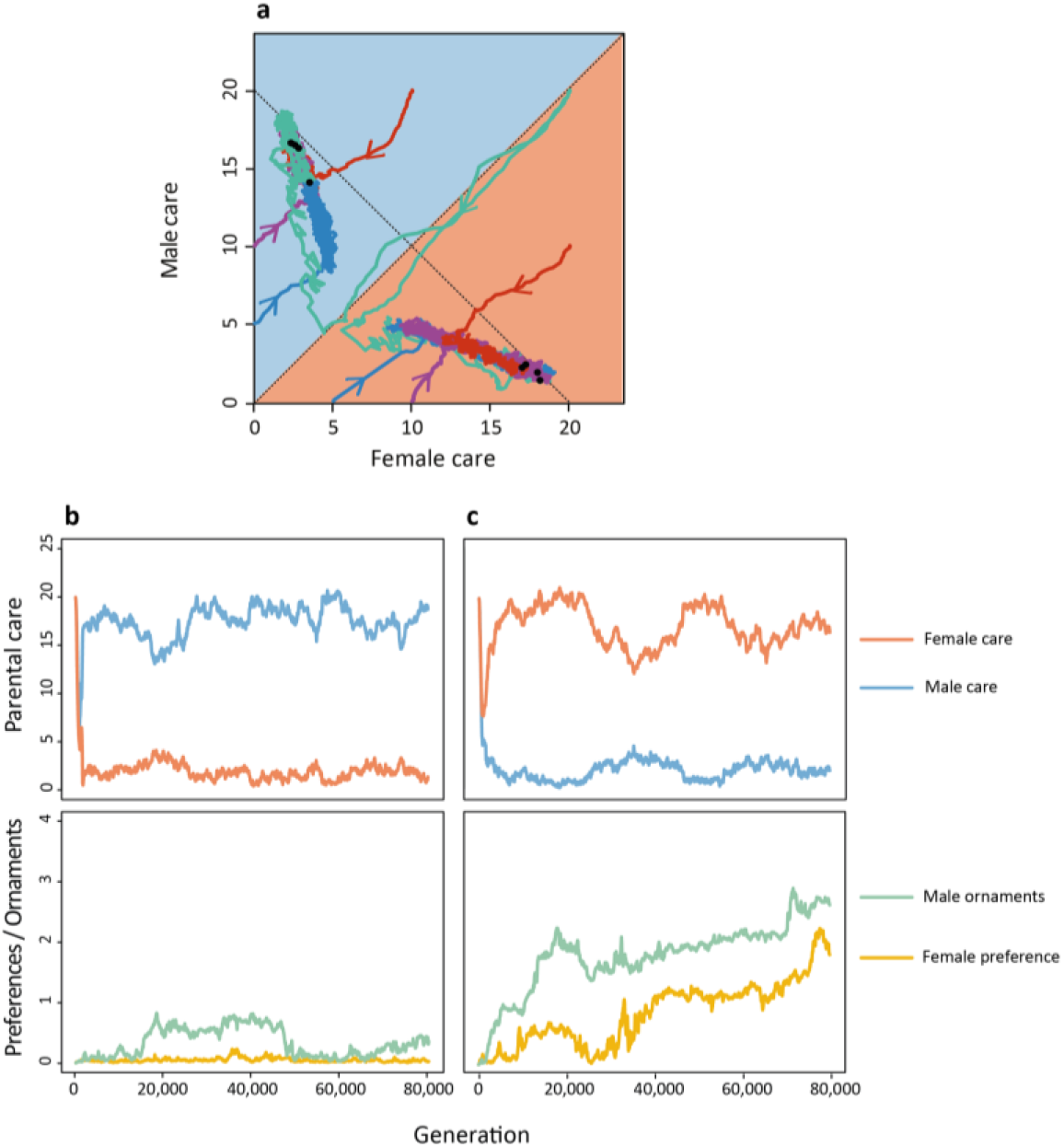
Joint evolution of mating and parental strategies. **(a)** If parental care strategies evolve alongside with the evolution of female preferences for a costly male ornament, all simulations result in one of two alternative equilibria. **(b)** One equilibrium is characterized by male-biased care, the absence of female preferences, and a small degree of male ornamentation. **(c)** The other equilibrium is characterized by female-biased care, strong female preferences, and a high degree of male ornamentation. In this simulation, there was no pre-mating period and no parental synergy.

Also these two types of equilibrium do not persist forever. As shown in Supplementary Fig. 4, each equilibrium defines the dominant sex role pattern for long periods of time (many thousands of generations), followed by a rapid switch to the other type of equilibrium. These transitions proceed in both directions. We investigated many of these transitions, and in all cases the parental strategy changed first (either from male-biased care to female-biased care, or *vice versa*), followed by the emergence or disappearance of female choosiness and male ornamentation. From this we tacitly conclude that, at least for the mating strategies considered in our simple model, the causal relationship goes from parental sex roles to mating roles, and not the other way around.

## Discussion

Here we investigated an individual-based simulation implementation of a modelling framework^25^ that may be viewed as the cornerstone of sex-role evolution theory. Although we made very similar assumptions as the analytical models, we arrived at remarkably different conclusions than the earlier mathematical analyses. First, the populations in our ‘null model’ (random mating, no sex differences in life-history parameters) do not evolve to egalitarian care^25^ or to a line (or curve) of equilibria^26^ but rather to one of two stable equilibria corresponding to strongly male-biased or strongly female-biased care, respectively. Second, our simulations suggest that even a small sex difference in pre-mating investment (like anisogamy) can induce the ‘Trivers effect’^12^ that the sex with the highest pre-mating investment is predestined for doing most of the post-mating parental care. This does not depend on factors as sexual selection or uncertainty of paternity, which can be expected to strengthen the Trivers effect. Third, parental synergy does not necessarily lead to egalitarian care. Even if it does, the evolutionary outcome is not necessarily efficient: in the presence of synergy the parents can be kept in a parental cooperation dilemma that in the absence of synergy is resolved by parental specialisation. Fourth, our simulations reveal that, as in the analytical models^25,26^sexual selection can lead to a situation where males are highly competitive on the mating market, while females provide most of the parental care. However, this is not the only outcome: there is a second equilibrium (that is equally likely) where males do most of the caring while the evolution of female choosiness is suppressed. Our simulations provide evidence that, in our model, the parental care pattern drives sexual selection and not the other way around^12^. Lastly, our simulations suggest that (parental and mating) sex roles are evolutionarily labile. For most of the parameters considered, the model has two stable equilibria.

Whenever this is the case, a simulation attains one of these equilibria for a long but limited period of time, followed by a rapid transition to the other equilibrium. Hence, male-biased care can switch to female-biased care, and *vice versa*. Similarly, a population can rapidly switch from a state of female choosiness, male competitiveness, and female-biased care to a state of male-biased care in the absence of choosiness and competiveness. These transitions occur for the same parameter settings; in contrast to other models (e.g. ref 55) they are not necessarily induced by a change in environmental conditions.

Why do our simulations lead to contrasting conclusions from the earlier analyses of very similar models? We think that our results highlight three limitations of analytical approaches that are mainly based on fitness considerations. As shown by Kokko & Jennions^25^ and Fromhage & Jennions^26^ the analysis of selection differentials and selection gradients can be very informative: they clearly indicate the effects of strategic parameters (like parental effort) on life history parameters (like own survival and offspring survival), thus quantifying the trade-offs between fitness components. However, selection-gradient based plots like Fig. 1a should not be over-interpreted, because it is not self-evident that evolution by natural selection proceeds in the direction of the selection gradient (the direction of steepest ascent of the fitness landscape). This only happens under restrictive assumptions, such as weak selection^56^, simple interactions across loci^57^, uncorrelated mutations of similar effect sizes^58^, and a simple structure of the genetic variance-covariance matrix^59^. A comparison of Fig. 1a and 1b shows that the gradient method predicts the simulation trajectories reasonably well when the fitness gradient is steep, but that it fails to detect directional selection away from egalitarian care when the curve of equilibria is approached (where the fitness gradient is close to zero). One could argue that the discrepancy between Fig. 1a and 1b is not too surprising, because a curve of equilibria, as predicted by the analytical model, is structurally unstable^60^ meaning that it will disappear if the model is slightly changes. However, we observed similar discrepancies in the parental synergy scenario where the gradient method predicts a structurally stable pattern of egalitarian care while the simulation model predicts the coexistence of two stable equilibria corresponding to either strongly male-biased or strongly female-biased care.

A second limitation of selection gradient methods is their focus on population averages. Averages have only a clear biological meaning if variation around them is small and symmetrically distributed^61^. In recent years, it is becoming increasingly clear that in the behavioural domain this assumption is not satisfied: in virtually all animals studied, individuals differ strongly and systematically in all kinds of behavioural tendencies^27,28,29^ (including parental^30,31,32^ and mating behaviour^62,63^), exhibiting so-called ‘animal personalities’^64^. Fig. 2 and 5a show that such individual variation in parental strategies, within and between the sexes, is also to be expected in the evolution of sex roles; in fact, it is shaped by natural selection (Supplementary Fig. 2). It has been argued before^35,36^ that such ‘patterned’ variation can strongly affect the course and outcome of evolution. This is clearly exemplified by our model, where the emergence of a bimodal distribution of care strategies is, in virtually all our simulations, the first step toward the evolution of sex role specialisation. The take-home message is that ‘selection gradient dynamics’ have to be interpreted with care if the emergence of individual variation is to be expected.

A third limitation of selection gradient approaches is their difficulty to include stochasticity. This is exemplified by our simulations including a pre-mating period (Fig. 3), where a rather subtle effect – the higher variance in mating success in the sex with the shorter pre-mating period, even in case of random mating – has a strong effect on the evolutionary outcome, providing a new underpinning for the Trivers effect.

At present, individual-based simulations are not yet very popular in evolutionary studies, presumably because of the belief that they do not add much to the evolutionary theory toolbox. Our study demonstrates that such simulations can be a useful check of analytical results, in particular in cases where the complexity of the evolutionary dynamics necessitates the usage of ‘short-cut’ methods (such as the selection-gradient method). On top of this, individual-based simulations have other advantages. They are easy to implement, without the necessity of performing complicated fitness calculations. For example, the fact that in the simulations each offspring has one mother and one father automatically guarantees that the ‘Fisher condition’ (that total reproductive success of all females is equal to the total reproductive success of all males) is satisfied, while the incorporation of this constraint in analytical models is not obvious^14,25,65,66^. Stochasticity, spatial structure, and environmental variation can easily be included in simulation models, in a variety of ways. The life cycle of the individuals can be much more intricate (and realistic) than in analytical models. Perhaps most importantly, individual interactions can be implemented in a natural way^37^. We have demonstrated how the evolution of mate choice can be included in the model, instead of representing sexual selection by constant parameters. This is relevant, because mating strategies and parental strategies must be allowed to evolve side by side in order to study evolutionary feedbacks between them. We are aware that our model of sexual selection is quite simple, but it is straightforward to include ‘good genes’ and ‘direct benefits’ variants^67,68^, as well as condition-dependent preferences^69^ and ornaments^70^.

We do not plead for replacing analytical methods by simulations. Simulations have the big disadvantage that their outcome can easily be ‘as complicated as reality’, thereby not furthering our understanding and sharpening our intuition. Instead, we recommend a pluralistic approach^71^ where analytical insights are checked and expanded by individual-based simulations, while the simulation outcomes are scrutinized with the help of analytical tools (such as the pairwise invasibility plots in Supplementary Fig. 2 and 7). The hope is to achieve a deeper understanding by a combination of diverse methods, in the spirit of Richard Levins’ insight^72^ (in our own wording): every model is a lie – all we can hope for is to approach truth by the intersection of independent lies.

## Methods

### Model structure

In line with the models of Kokko and Jennions^25^ and Fromhage and Jennions^26^, we consider a population with overlapping generations and discrete time structure. To be concrete, we assume that a time unit corresponds to one day. The population consist of females and males that, on each day, can be in one of the following states: juvenile, pre-mating, mating, or caring. In each of the four states, there is a fixed mortality rate, which can be sex-specific. Unless stated otherwise, all mortalities were set to 0.001 day^−1^. Therefore, the expected lifespan of an individual is 1000 days, a value that we consider a proxy for generation time. Offspring mortality is density dependent, thus ensuring a limited population size. In our baseline scenario, population size fluctuates around 2000 females and 2000 males.

The life cycle of our model organisms is illustrated in Supplementary Fig. 1. Offspring that survive the period of parental care spend a fixed number of days (the maturation time) in the juvenile state. In all simulations reported, the maturation time of both sexes was equal to 20 days. After maturation, the surviving individuals enter the pre-mating state, corresponding to a condition where they prepare for mating (e.g. territory establishment; nest building; replenishment of gametes). After a fixed sex-specific number of days, the pre-mating state changes into the mating state. Unless stated otherwise, the pre-mating period was set to zero, meaning that individuals move to the mating state without delay. Once in the mating state, individuals seek for mating opportunities. In our baseline scenario, females and males mate at random, but we also consider a mate-choice scenario where females have a preference for certain male ornaments. On a given day, mating is modelled as follows: one by one, a female in the mating state is selected at random. As long as there are still males in the mating state, the female encounters one of these males at random. In the random mating scenario, such an encounter always results in mating; in the mate-choice scenario, the male can be rejected if its ornamentation does not fit to the preference of the female (see below). When mating does occur, both the male and the female immediately leave the mating state and both enter the caring state. When a female-male encounter does not result in mating, both individuals stay in the mating state, but they are no longer available for mating on that day. Hence each individual in the mating state can only have one encounter per day, and a female and a male both lose one day if their encounter does not result in mating. Mating will stop for the day when no more males in mating state are available and/or when all females in mating state have made their mating decisions. All remaining individuals stay in the mating state, but they will only have a new mating opportunity on the following day.

Once a mating has occurred, the mated couple produces a clutch of offspring. Offspring survival strongly depends on the amount of parental care received. The female care duration *T*_*f*_ and the male care duration *T*_*m*_ are heritable traits that may differ between individuals. The evolution of *T*_*f*_ and *T*_*m*_ is the core subject of our study. We interpret *T*_*f*_ and *T*_*m*_ as the ‘intended’ cared duration: if one of the parents dies during the care period, this intended care duration is replaced by the actual care duration (the time from mating to death). To consider the possibility of synergy between the two parents, we assume that their total parental effort is given by *T*_*tot*_ = *T*_*f*_ + *T*_*m*_ + *σT*_*f*_*T*_*m*_ where the ‘synergy’ parameter *σ* is non-negative. Unless stated otherwise, we assume that *σ* = 0, meaning that each parent has an independent additive effect on total care. Offspring survival is proportional to 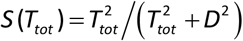 an increaseng sigmoidal function of total parental care. The parameter *D* may be viewed as a measure of the care demand of offspring: the function *S* has a turning point at *T*_*tot*_ = *D*, implying that the marginal benefits of care are maximal when the total parental effort matches *D*. Throughout, we consider the case *D* = 20, i.e. the offspring demand the equivalent of 20 days of care. When the care period *T*_*f*_ (resp. *T*_*m*_) has passed, the corresponding parent changes into the pre-mating state. When the longest-caring parent stops caring, the surviving offspring enter the infant state. As mentioned above, population size is regulated in our model by assuming that offspring survival is density dependent: it is given by *S* (*T*_*tot*_)/(1 + *γN*), where *N* is the current population size and the parameter γ quantifies the degree of density dependence. This form of density regulation ensures that expected lifetime reproductive success (the fitness measure used by analytical approaches; see below) does indeed predict the course and outcome of evolution^73^. Our choice *γ* = 0.003 ensured relatively large populations (about 2000 females and 2000 males) with limited genetic drift and demographic stochasticity.

At the start of a new day, the survival of each individual was checked according to the individual’s sex- and state-specific mortality. Non-survivors were removed from the population.

### Sexual selection

In part of our study, we consider a mate-choice scenario where females can evolve a preference *p* for a male trait of size *s*, where *p* and *s* are both heritable traits. In line with Kokko and Johnstone^52^, we assume that the probability that a female with preference *p* that encounters a male with trait size *s* will actually mate with this male is given by the logistic expression (1 + *κ* exp(*α*(*p* − *s*)))^−1^. For all non-negative values of *p*, this expression increases with *s* (hence all females have a preference for males with larger ornament sizes), and the rate of increase is positively related to *p* (hence females with a large value of *p* discriminate more strongly against males with a small trait size). The parameters *κ* and *α* are scaling factors that affect the intensity of sexual selection. The mate-choice simulations shown are all based on the parameter values *κ* = 0.02 and *α* = 2. For these parameters, an ‘unattractive’ male with *s* = 0 is accepted for mating with probability 0.98 by a female with a preference value is almost undistinguishable from random mating) and with probability 0.48 by a female with preference value *p* = 2. We assume that male ornamentation is costly: each time step, the survival probability of a male with trait size *s* is reduced by a percentage *βs*^2^, where we chose β = 10^−6^.

### Reproduction and inheritance

For simplicity, we consider a population of haploid individuals that may differ in their alleles at four gene loci. The *T*_*f*_-locus and the *p*-locus are only expressed in females, and the *T*_*m*_-locus and the *s*-locus are only expressed in males. The alleles at the *T*_*f*_-locus and the *T*_*m*_-locus determine the duration of maternal and paternal care, respectively. The allele at the *p*-locus determines the degree of female preference, while the allele at the *s*-locus determines the size of the male trait. In our baseline scenario (random mating), the *p*-allele and the s-allele are not expressed. Offspring inherit their alleles from their parents’ subject to mutation. In a first step, the allele at each locus is drawn at random from one of its parents. Moreover, offspring sex is determined at random, with equal probability. In a second step, mutations could occur with probability *μ* = 0.005 per locus. If a mutation occurs at the *T*_*f*_-locus or the *T*_*m*_-locus, the current allele is either increased or decreased by 1, with equal probability. This ensures that the parental care times *T*_*f*_ and *T*_*m*_ are natural numbers. If a mutation occurs at one of the other two loci, a small mutational step of size ɛ was drawn from a Cauchy distribution (with location parameter 0 and scale parameter 0.01) and added to the current value of *p* or *s*, respectively. We used the Cauchy distribution (rather than a normal distribution) because it allows for occasional larger step sizes. However, we limited mutational step sizes to a maximum value of *ɛ*_max_ = 0.05.

### Initialization and replication

In all simulations, the *p*- and the *s*-locus were initialized at *p* = *s* = 0. The *T*_*f*_-locus and the *T*_*m*_-locus were initialized at different values (leading to the different trajectories in Fig. 1b and 6a); each time, we started with a monomorphic population. For each parameter combination, we ran at least 100 replicate simulations. In all cases, the outcome was highly repeatable, allowing us to focus on one or two replicates. As partly documented in the Supplement, we also ran numerous simulations for model variants that differed from the baseline model in its parameter values (state- and sex-specific mortalities; offspring demand *D*; cost of ornamentation *β*; density dependence *γ*; mutation rate *μ*), the survival function *S* (*T*_*tot*_), the mate choice function, or the distribution of mutational step sizes. In all cases, we arrived at the same conclusions as reported in the manuscript. We therefore conclude that our results and conclusions are quite robust.

### Mathematical analysis

As a standard of comparison for our individual-based simulations, Fig. 1a shows the trajectories of the corresponding deterministic model, making use of the fitness gradient method described in Kokko and Jennions^25^ and Fromhage and Jennions^26^. In a nutshell, this method calculates the selection gradient (indicating the strength and direction of selection) in males and females for each combination of parental care parameters (*T*_*f*_,*T*_*m*_). This gradient points into the direction of steepest ascend of the fitness landscape, where fitness is defined by expected lifetime reproductive success. Under the assumption that evolution will proceed in the direction of the selection gradient, evolutionary trajectories as in Fig. 1a are obtained. Our model is inspired by the model of Kokko and Jennions^25^ and Fromhage and Jennions^26^, but it differs from the former models in various respects. In the Supplement, we discuss these differences and demonstrate that our main results are also recovered for the earlier models, again indicating the robustness of our results and conclusions.

## Data availability

This study is theoretical; no new empirical data were generated.

## Code availability

The C++ simulation code and a Mathematica file with an implementation of the fitness gradient method are available for download from https://github.com/xiaoyanlong/evolution-of-sex-roles.

## Acknowledgements

We thank G.S. van Doorn, R. Scherrer, J. Komdeur, T. Székely and the MARM group at the University of Groningen for valuable discussion, comments, and suggestions. We are grateful to L. Fromhage for sharing technical details of the fitness gradient method and some computational resources with us. We appreciate H. Hildenbrandt and M. Mosna helped with our programming. We would like to thank the Center for Information Technology of the University of Groningen for their support and for providing access to the Peregrine high performance computing cluster. X.L. is supported by a PhD fellowship of the Chinese Scholarship Council (NO. 201606380125). F.J.W. acknowledges funding from the European Research Council (ERC Advanced Grant No. 789240).

## Author contributions

Both authors conceived the study and developed the model. X.L. performed the computational work and analysed the data. Both authors interpreted the results and wrote the manuscript.

