## Supplementary information for "Individual variation in parental care drives divergence of sex roles"

Supplementary Figure 1: Diagram illustrating the life cycle in our model.

Supplementary Figure 2: Evolutionary branching of parental care strategies.

Supplementary Figure 3: Effect of population size on the timing of evolutionary transitions.

Supplementary Figure 4: Transitions between alternative mating and caring strategies

Supplementary Figure 5: Rescaled version of the Fromhage & Jennions (2016) model.

Supplementary Figure 6: Evolution of sex-biased care in the rescaled FJ model.

Supplementary Figure 7: Pairwise Invasibility Plot of the rescaled FJ model.

Supplementary References

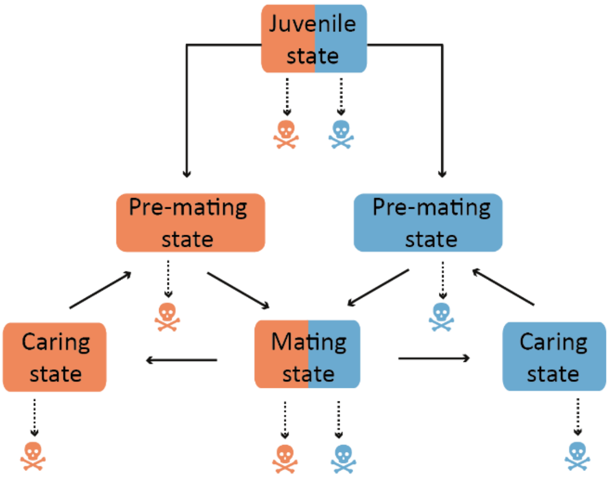


**Supplementary Fig. 1 | Diagram illustrating the life cycle in our model.** Offspring surviving the parental-care period enter the 'juvenile state' where they stay for a fixed (and potentially sex-specific) maturation time. Afterwards, they spend a fixed time period (which is zero in the baseline version of the model) in the 'pre-mating state'. Then they enter the 'mating state' where they randomly encounter individuals of the other sex. In case of mate choice, not every encounter results in a mating. If a mating does occur, both mating partners switch to the 'caring state', where they stay for a genetically determined time period ($T_f$ or $T_m$, respectively). The total care duration has a positive effect on the survival of their offspring. Once the care period of a parent has been completed, the individual switches to the pre-mating state, from where the whole cycle repeats itself. In all states, mortality can occur (which potentially is sex-specific). In the baseline version of the model, individual life expectancy is 1000 time units (= 'days'). For simplicity, we equate this time period with one 'generation'. Colour conventions: throughout the manuscript females are indicated by the colour red, and males are indicated by the colour blue.

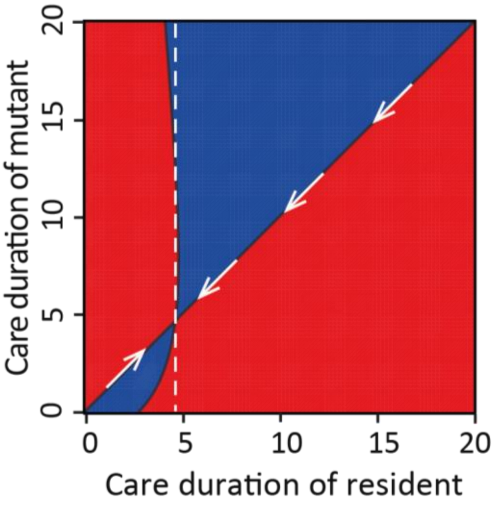


**Supplementary Fig. 2 | Evolutionary branching of parental care strategies.** The graph shows a 'Pairwise Invasibility Plot (PIP)' of the baseline version of our model (random mating, no differences between the sexes). We restrict attention to egalitarian care ($T_f = T_m$), allowing us to conduct a one-dimensional analysis. A PIP illustrates which mutant strategies can invade when rare in a given resident population. As explained in detail in Geritz et al. (1998), the *x*-axis depicts all possible care durations of a resident population, while the care durations of mutants are represented on the *y*-axis. The red area of the plot corresponds to mutant-resident combinations where the mutant has a higher fitness than the resident and, hence, can invade the resident population. Here, fitness is calculated as in Fromhage & Jennions (2016). The blue area indicates those mutant-resident combinations where the mutant has a lower fitness than the resident and is selected against. The separating black lines corresponds to situations where mutants and residents have the same fitness. The two equal-fitness lines intersect at the value $T^* = 4.67$, which is a so-called 'Evolutionarily Singular Strategy'. The white arrows indicate that this strategy is 'convergence stable': in the course of evolution, the resident population is shifted toward $T^*$. The dashed vertical line lies in the red region (at least for mutants close to $T^*$), implying that there are mutants that can invade the resident $T^*$. This means that $T^*$ is *not* evolutionarily stable. A configuration like this (convergence to an evolutionarily unstable strategy) is called a 'branching point', because it indicates that directional selection (toward $T^*$) switches to disruptive selection (once $T^*$ is reached) indicates that directional selection (toward $T^*$) switches to disruptive selection (once $T^*$ is reached). When the population would remain constrained to egalitarian care, a dimorphic population would result, where part of the population would care less than $T^* = 4.67$ while another part would care more than this value. If sex differentiation in care is possible, it is to be expected that each of the 'branches' gets associated with one of the two sexes (e.g. low-care might get associated with the female sex and high-care with the male sex) (see Rueffler et al. 2006). A *Mathematica* file with the implementation of the PIP is available via the link in the main text.

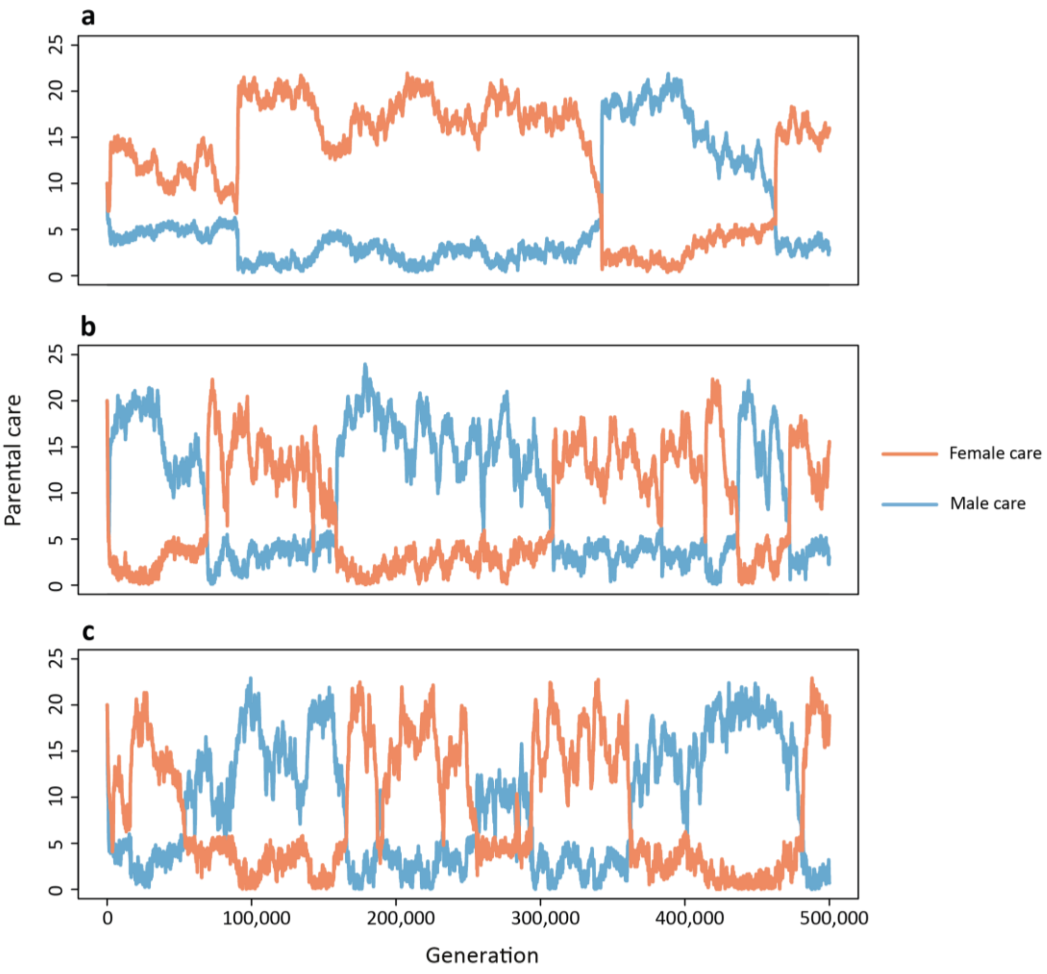


**Supplementary Fig. 3 | Effect of population size on the timing of evolutionary transitions.** As shown in Fig. 4 of the main text, parental roles are evolutionarily labile in that spontaneous transitions occur from one parental-care equilibrium to the other. As explained in the main text, the mean time between transitions depends on the duration of the pre-mating period (affecting the strength of selection) and on population size (affecting genetic drift). The three panels illustrate, for simulations without pre-mating period, how the number of transitions within a fixed period of 500,000 generations increases with a decrease of population size. **(a)** 2,000 females and 2,000 males: two transitions; **(b)** 650 males and 650 males: five transitions; **(c)** 300 females and 300 males: six transitions. Population sizes fluctuated and were regulated by changing model parameter $\gamma$ (see Methods).

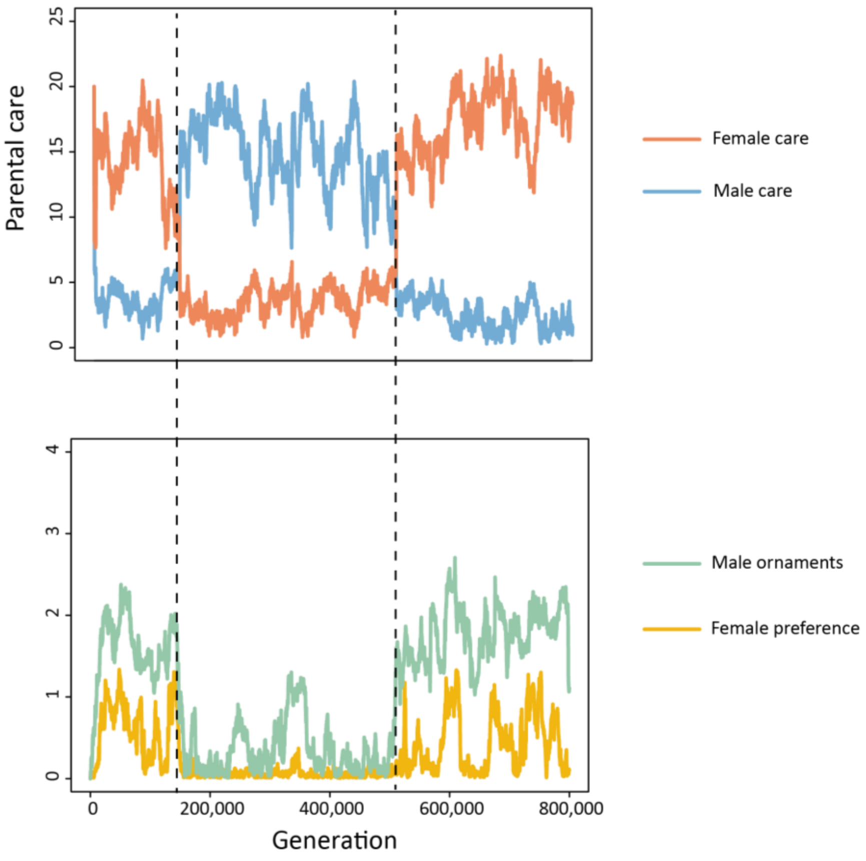


**Supplementary Fig. 4 | Transitions between alternative mating and caring strategies.** As shown in Fig. 6 of the main text, the joint evolution of mating and parental care strategies leads to one of two equilibria: male care in the absence of female choosiness and female care, female care associated with a female preference for ornamented males and costly male ornamentation. The simulation demonstrates that also these combined mating and parental roles are evolutionarily labile in that spontaneous transitions occur from one equilibrium to the other. Epochs with female choosiness and female-biased care (here: first 180,000 generations, last 300,000 generations) alternate with epochs with random mating (= no female preference) and male-biased care (here: generations 180,000 till 500,000). A more detailed analysis revealed that the change in female preferences was always preceded by a change in the parental care strategies of the two sexes.

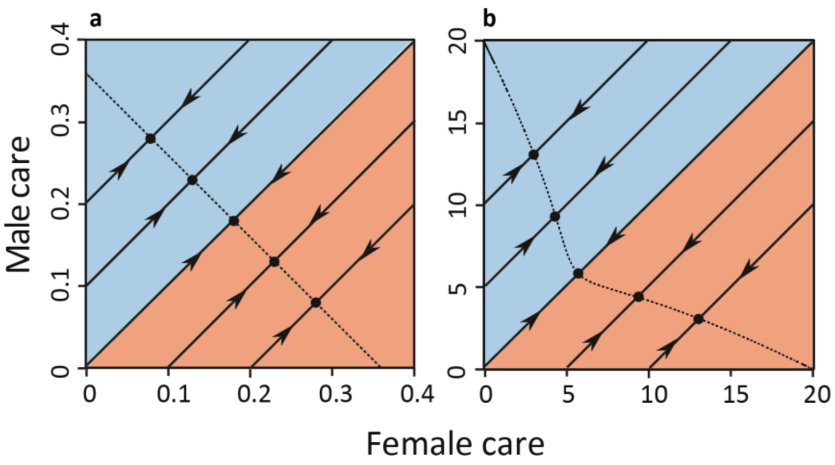


**Supplementary Fig. 5 | Rescaled version of the Fromhage & Jennions (2016) model.** The model presented here was inspired by the 'FJ model' of Fromhage & Jennions (2016). However, we replaced their offspring survival function $S(T_{tot}) = \exp(-D/T_{tot})$ by the more symmetric function $S(T_{tot}) = T_{tot}^2 \big/ \left(T_{tot}^2 + D^2\right)$. **(a)** For their parameter setting $D = 0.1$ and $\mu = 0.01$ (where $\mu$ denotes the mortality rate per day), Fromhage and Jennions concluded that the selection gradient method predicts convergence to a line of neutrally stable equilibria (Fig. 1a of their article). **(b)** For the parameters used in our model ($D = 20$ and $\mu = 0.001$), the FJ model produces an almost identical pattern as our variant of the model (see Fig. 1a in our main text). This shows that, also in the FJ model, the 'curve of equilibria' is not necessarily a straight line.

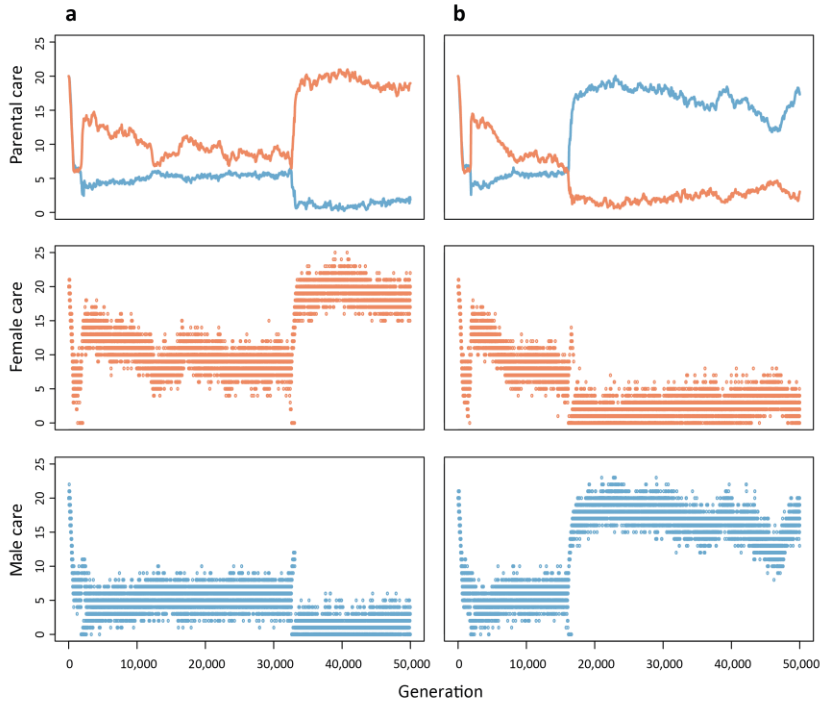


**Supplementary Fig. 6 | Evolution of sex-biased care in the rescaled FJ model.** The rescaled version of the FJ model ($D = 20$ and $\mu = 0.001$, see Supplementary Figure S6(b)) exhibits a very similar behaviour as our version of the model. Two representative simulations show **(a)** the evolution of female-biased care; and **(b)** the evolution of male-biased care. On a long-term perspective, transitions between the two types of equilibria also occurred. Notice that a longer period of low-level egalitarian care precedes the first switch to sex-biased care. This is explained by the Pairwise Invasibility Plot in Supplementary Fig. 7.

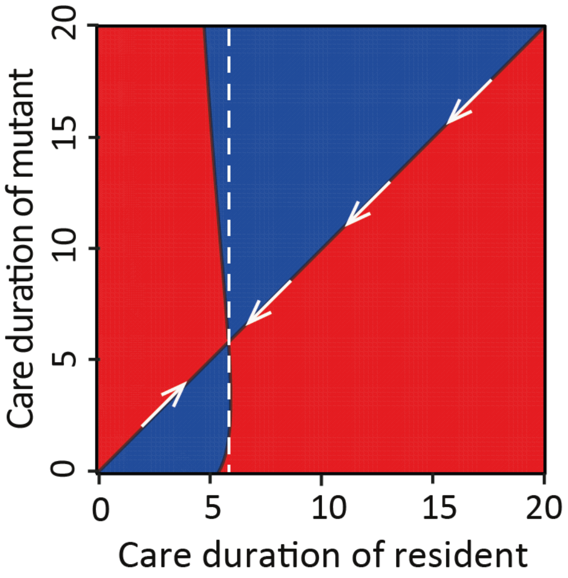


**Supplementary Fig. 7 | Pairwise Invasibility Plot of the rescaled FJ model.** With the same method as in Supplementary Fig. 2, we constructed a PIP for the rescaled FJ model (parameters $D = 20$ and $\mu = 0.001$, see Fig. S6(b)). Now there is a convergence stable singular strategy at $T^* = 5.87$. In contrast to our version of the model (Supplementary Figure S2), this singular strategy is not a branching point but evolutionarily stable (mutants close to $T^*$ cannot invade, as the dashed vertical line lies in the blue for mutants close to $T^*$). Standard adaptive dynamics theory (Geritz et al. 1998) would therefore predict that egalitarian care at level $T^*$ is an evolutionary attractor, and, hence, and endpoint of evolution. In contrast, *all* simulations resulted in the evolution of sex-biased care (see Supplementary Fig. 6). Similar observations were made in other simulation studies (Wolf et al. 2007, Berngruber et al. 2010, Baldauf et al. 2012), where diversification occurred at an evolutionary attractor. In all these cases, $T^*$ is locally but not globally evolutionarily stable, as the dashed vertical line transverses the red region as well (mutants with a very short care duration can invade the population of $T^*$ residents. If one waits long enough, such mutants will invariably appear in individual-based simulations (and in the real world). As argued in Wolf et al. (2008), the analytical conditions for evolutionary branching are based on the assumption of infinitesimally small mutational step sizes and therefore correspond to a 'worst-case scenario for evolutionary diversification'. In simulations, diversification (or, as in our case, sex differentiation) can predictably occur under much milder conditions.
